# Organometallic gold(III) [Au(Hdamp)(L1^4^)]Cl (L1 = *SNS*-donating thiosemicarbazone) complex protects mice against acute *T. cruzi* infection

**DOI:** 10.1101/312702

**Authors:** Carla Duque Lopes, Ana Paula S. Gaspari, Ronaldo J. Oliveira, Ulrich Abram, José P. A. Almeida, Pedro v. S. Maia, João S. da Silva, Sérgio de Albuquerque, Zumira A. Carneiro

**Affiliations:** Department of Clinical Toxicological and Bromatological Analysis School of Pharmaceutical Sciences of Ribeirão Preto, University of São Paulo (USP), Ribeirão Preto, Brazil; Núcleo de Desenvolvimento de Compostos Bioativos (NDCBio), Universidade Federal do Triângulo Mineiro, Uberaba – MG, Brazil; Freie Universität Berlin, Institute of Chemistry and Biochemistry, Berlin, Germany; Centro Universitário Estácio de Ribeirão Preto, SP, Brazil; Departament of Biochemistry and Immunology, School of Medicine, University of São Paulo, Ribeirão Preto, São Paulo, Brazil; Departamento de Química, Faculdade de Filosofia, Ciências e Letras de Ribeirão Preto – FFCLRP-USP, University of São Paulo, Ribeirão Preto, SP, Brazil

**Author notes:** These authors contributed equally. Address correspondence to Zumira Carneira.

**Keywords:** Gold(III) Complex, Thiosemicarbazones, Chagas disease, Trypanocidal activity, immune response, IFN-γ production

## Abstract

Chagas disease remains a serious public health concern with unsatisfactory treatment outcomes due to strain-specific drug resistance and various side effects. To identify new therapeutic drugs against *Trypanosoma cruzi*, we evaluated both the *in vitro* and *in vivo* activity of the organometallic gold(III) complex [Au(Hdamp)(L1^4^)]Cl (L1 = *SNS*- donating thiosemicarbazone), which was denoted 4-Cl. Our results demonstrated that 4- Cl was more effective than benznidazole (Bz) in eliminating both the extracellular trypomastigote and the intracellular amastigote forms of the parasite without cytotoxic effects on mammalian cells. In very-low-dose *in vivo* assays, 4-Cl reduced parasitaemia and tissue parasitism in addition to protecting the liver and heart from tissue damage. All these changes resulted in the survival of 100% of the mice treated with 4-Cl during the acute phase. We hypothesised that 4-Cl can act directly on the parasite and may participate in the modulation of IFN-γ production at the acute stage of the disease. Molecular docking simulations showed that the compound may interact with cruzain, a thiol protease considered a possible antiparasitic drug target, primarily by hydrophobic interactions. These analyses predicted that the Cys25 residue in the cruzain binding site is approximately 3.0 Å away from the S and Au atoms of the gold compound, which could suggest formation of a possible covalent bond between cruzain and the inhibitor. Overall, we confirmed the potential of 4-Cl as a new candidate for Chagas disease treatment.

## Introduction

Chagas disease is a neglected infection caused by a protozoan parasite named *Trypanosoma cruzi (T. cruzi)*, transmitted by triatomine insect vectors, which is endemic in Latin America. According to the World Health Organization (WHO), approximately ten million people are infected with *T. cruzi* worldwide(1, 2). In addition, the phenomena of globalization and immigration have also led to the appearance of several infectious cases in development of the disease(2). Currently, only two drugs are available for the treatment of the Chagas disease: nifurtimox (NFX) and benznidazole (Bz)(3, 4). In some countries, NFX was discontinued due to serious side effects, such as neuropathy and anorexia, among others(5, 6). Thus, the only drug currently used for therapeutic purposes is Bz which is effective only during the acute phase of the infection but may present undesirable systemic toxicity, such as rashes and gastrointestinal symptoms(6). Therefore, the development of more efficacious and less toxic drugs, which can be used as alternatives for drug resistance, is urgently needed(7, 8).

A potential strategy for the treatment of Chagas disease is the design of compounds that selectively inhibit essential enzymes for parasite survival inside the host cells(9). In the case of *T. cruzi*, cruzain is an attractive drug target. The structure of cruzain contains a cysteine protease domain, which plays an important role during the life cycle of the parasite, such as replication, metabolism, and evasion of host immune defence during the early events of macrophage infection(10). Although many other potential drug targets exist in parasite metabolism(8), cruzain is by far the most studied protease of *T. cruzi* due to its role as a virulence factor of the parasite(9). Thus, compounds that inhibit the biological function of cruzain, such as thiosemicarbazones, may be an effective alternative for pharmacological treatment of Chagas disease(6, 7). In this context, transition metal complexes that have thiosemicarbazones as ligands have been developed and tested against various forms of *T. cruzi* (8, 9). Gold(III) complexes have received increased attention due to their biological properties, such as anti-cancer(11) and antiparasitic effects(12). In a recent paper, our research group identified complexes of the general formula [Au^III^(Hdamp)(L1)]Cl (Fig. 1) that have high stability in aqueous solution and antiparasitic activity against the *Tulahuen LacZ* strain. Among these compounds, 4-Cl, [Au^III^(Hdamp)(R_1_R_2_L1^4^)]Cl (R_1_ = R_2_ = Methyl) demonstrated low cytotoxicity in spleen cells, leading to a selectivity index (SI) of approximately 30(8), which indicated that this compound is a promising candidate for the development of trypanocidal drugs.

**Fig 1.**
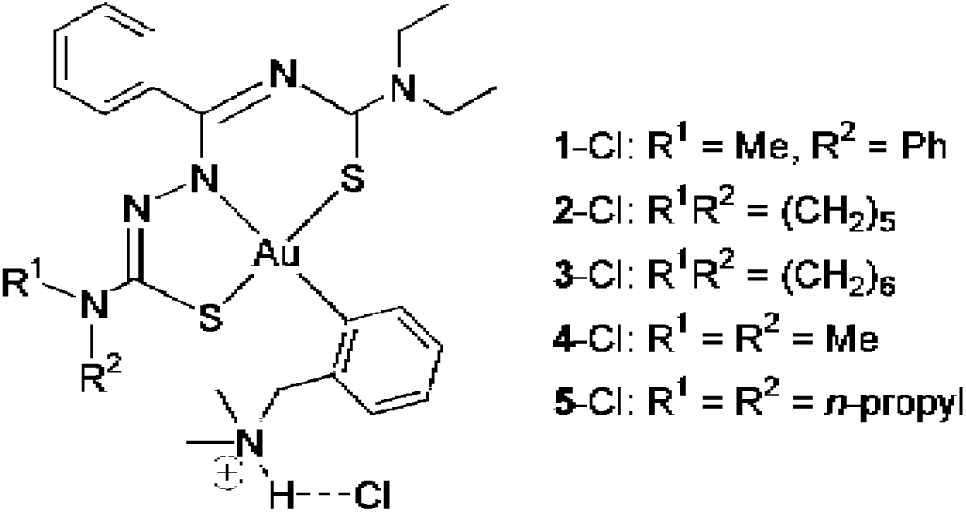
Organometallic gold(III) complexes containing hybrid *SNS*-donating thiosemicarbazone ligands [Au^III^(Hdamp)(L1)]Cl (Hdamp = dimethylammoniummethylphenyl) (Adapted from (8)).

Clinical manifestations associated with *T. cruzi* infection are dependent on the intricate equilibrium between the parasite and the host immune response. The production of interferon gamma (IFN-γ) generated for adaptive immune responses is important for inhibition of parasite proliferation(13, 14). Intense production of IFN-γ activates the CD4 T lymphocytes with s Type 1 Helper T (Th1) profile and cytotoxic CD8 T lymphocytes, which efficiently eliminate the parasite (15). T-Helper 17 (Th17) cells have the potential to become antigen-specific CD8 T cells against the parasite and show greater contributions than the subpopulation of CD4 T cells, which are producers of IFN-γ. Consequently, their cytotoxic and cytokine secretion functions are decreased, hindering parasite elimination (16). Therefore, the modulation of immune responses is fundamental for a good prognosis of Chagas disease.

Since the acute phase of Chagas disease has been associated with *T. cruzi* II-restricted infections, in the present study, the *in vitro* and *in vivo* 4-Cl trypanocidal activity was evaluated against the Y strain (group TcII) to assess the use of this compound as a new drug for the treatment of Chagas disease. Docking studies were also conducted to elucidate the interaction between the gold(III) complex with the *T. cruzi* enzyme cruzain (18). Based on the cruzain inhibitory potential of thiosemicarbazone-derived compounds, molecular docking techniques were employed on this enzyme to identify the possible target of [Au(Hdamp)(L1^4^)]Cl in the parasite.

## Results

### The gold complex acts directly on *T. cruzi* parasites

The organometallic gold(III) complex 4-Cl has interesting characteristics from both chemical and biological points of view (8). Then, we initially assessed the effectiveness of 4-Cl against the *T. cruzi* Y strain, which is partially resistant to Bz treatment and more virulent than the *Tulahuen* strain. The trypomastigotes of the Y strain were incubated with serial dilutions of 4-Cl or Bz for 24 h, and live parasites were counted by colorimetric analyses. The 4-Cl treatment was highly efficient compared to Bz treatment, reaching an IC_50Try_ (concentration needed to kill 50% of the parasites) of 0.03±0.006 μM, whereas the IC_50Try_ of Bz was 0.96±0.025 μM; these results indicate that 4-CL is almost thirty-two times more effective than Bz in killing the trypomastigote forms of *T. cruzi* (Fig. 2A). To determine the effects of 4-Cl on intracellular amastigotes of the *T. cruzi* Y strain, BMMs were differentiated and infected with the trypomastigote forms for 16 h. The extracellular parasites were removed by extensive washes and incubated with low concentrations of 4-Cl or Bz for 24 hours. The Bz treatment maintained the same percentage of parasite killing at both concentrations, and its trypanocidal capacity decreased with subsequent concentrations. At 3.2 μM, 4-Cl killed 75% of the intracellular amastigotes of *T. cruzi* and was more efficient than Bz (54% inhibition of the replication or survival of the amastigotes). This high efficacy remained until a concentration of 1.56 μM, and its trypanocidal activity was reduced only at nanomolar concentrations (Fig. 2B). Interesting, the gold(III) complex at the highest concentrations lost its ability to eliminate the intracellular parasites. This phenomenon was not due to cytotoxicity in macrophages, whose the EC_50(4-Cl)_ was 70 μM, relatively close to the EC_50_ in primary culture of spleen cells (113 μM) (8). Most likely, at this high concentration, there is a saturation of the absorption of the compound, which is no longer effective in eliminating the parasite. The same phenomenon was observed for the highest concentration of Bz (Fig. 2B). Therefore, by excluding the values of 6.2 μM to obtain a real IC_50,_ for both compounds, we obtained an IC_50(ama)_ of 0.5 μM for 4-Cl, indicating a trypanocidal activity 3.6 times higher than that of Bz (IC_50(ama)_ of 1.8 μM). Together, these data indicate that 4-Cl is more efficient in killing both mammalian forms of the Y strain of *T. cruzi* than Bz.

**Fig 2.**
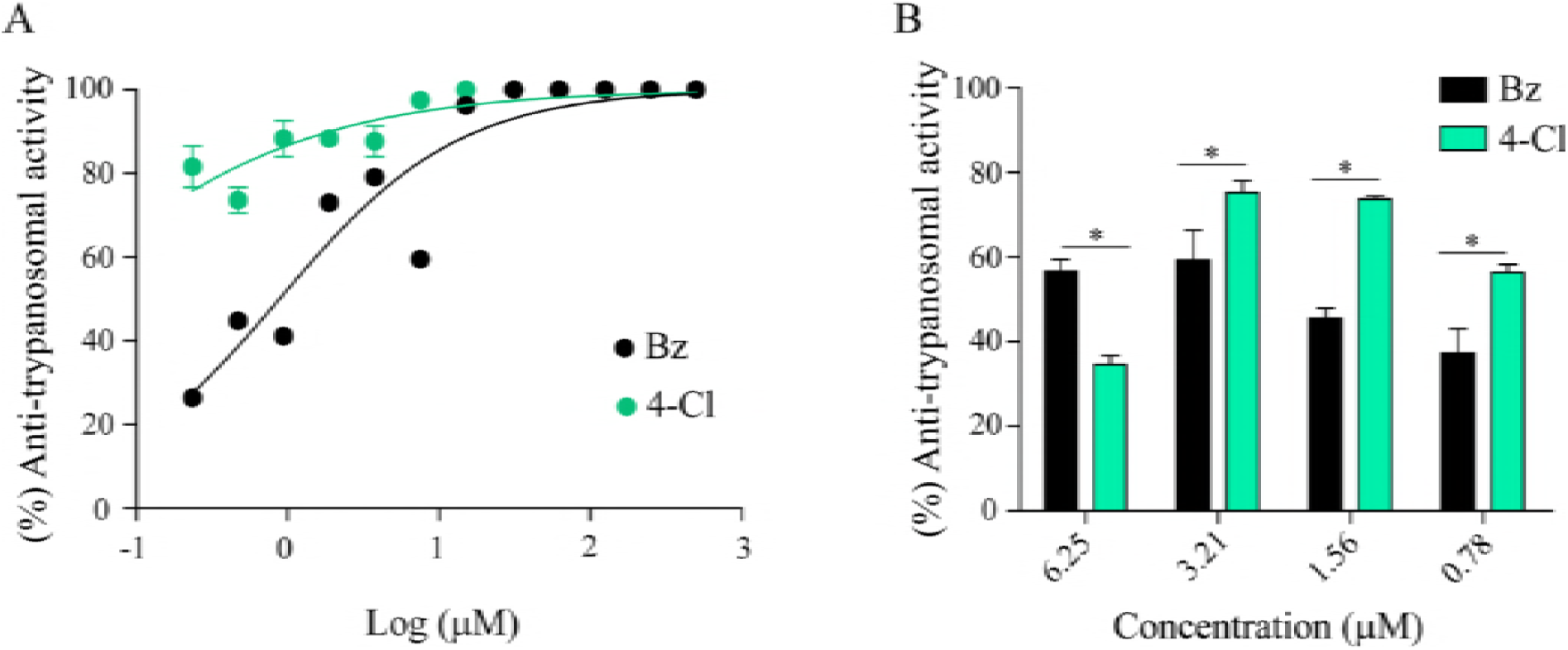
*In vitro* trypanocidal activity of 4-Cl against the trypomastigote and amastigote forms of the Y strain of *T. cruzi.* (A) Percentage of trypanocidal activity of 4-Cl and Bz against the Y strain of *T. cruzi* analysed by quantifying viable parasites 24 h post-treatment. (B) Macrophages derived from bone marrow were infected with the trypomastigote form of the Y strain of *T. cruzi*. After 24 hours of infection, the extracellular parasites were removed, and cells were infected with the amastigote forms and treated with 4-Cl or Bz. Assays were conducted using biological replicates.

### The 4-Cl treatment protects the mice from *T. cruzi* infection

The high efficiency of this treatment in eliminating both trypomastigote and amastigote forms *in vitro* prompted us to investigate the potential of 4-Cl *in vivo*. For these analyses, mice were infected with 2000 blood-derived trypomastigote forms of the Y strain and orally treated with 2.8 mg/kg/day of 4-Cl, two different concentrations of Bz, 2.8 mg/kg/day of Bz(-) or 100 mg/kg/day of Bz(+) for 10 consecutive days from the fifth day of infection (first point of parasitaemia). To optimise the treatment dose of 4- Cl, we administered different concentrations to the mice based on the clinical dose of Bz(+). At the highest doses, 4-Cl showed several toxicities and did not control the parasitaemia (Fig S1). However, at a dose of 2.8 mg/kg, 4-Cl eliminated 61.3% of the circulating parasites at the peak of infection (9 d.p.i.), while Bz(-) showed no reduction compared to that of the PBS group (Fig. 3A). To determine whether the blood reduction of parasites in 4-Cl-treated mice was reflected in the *T. cruzi* migration to skeletal tissues, we performed a qPCR analysis specific to *T. cruzi* DNA in the heart and skeletal muscle after 15 d.p.i. The number of copies found in the heart of 4-Cl-treated mice was similar to Bz(-) treatment but was significantly less than that of the PBS-treated group (Fig. 3B). In skeletal muscle, 4-Cl significantly reduced the parasitism compared with both Bz(-) and PBS, showing that a low dose of 4-Cl is effective at preventing the systemic spread of *T. cruzi* (Fig. 3C). Interestingly, evaluation of the heart histological sections showed that the number of nests of both 4-Cl- and Bz(-)-treated mice was reduced (Fig. 3D). Heart histological analysis revealed that amastigote nests (Fig. 3E-G) decreased in all treated groups compared to the controls (Figure 3E). The amastigote nests in the Bz(-) treatment group were the smallest and most scattered (Fig. 3F), in contrast to the findings for the 4-Cl group, which showed the largest and most individualised nests (Fig. 3G) which may justify that same amount of *T. cruzi* DNA was found in the Bz(-) and 4-Cl heart samples (Fig. 3B).

**Fig 3.**
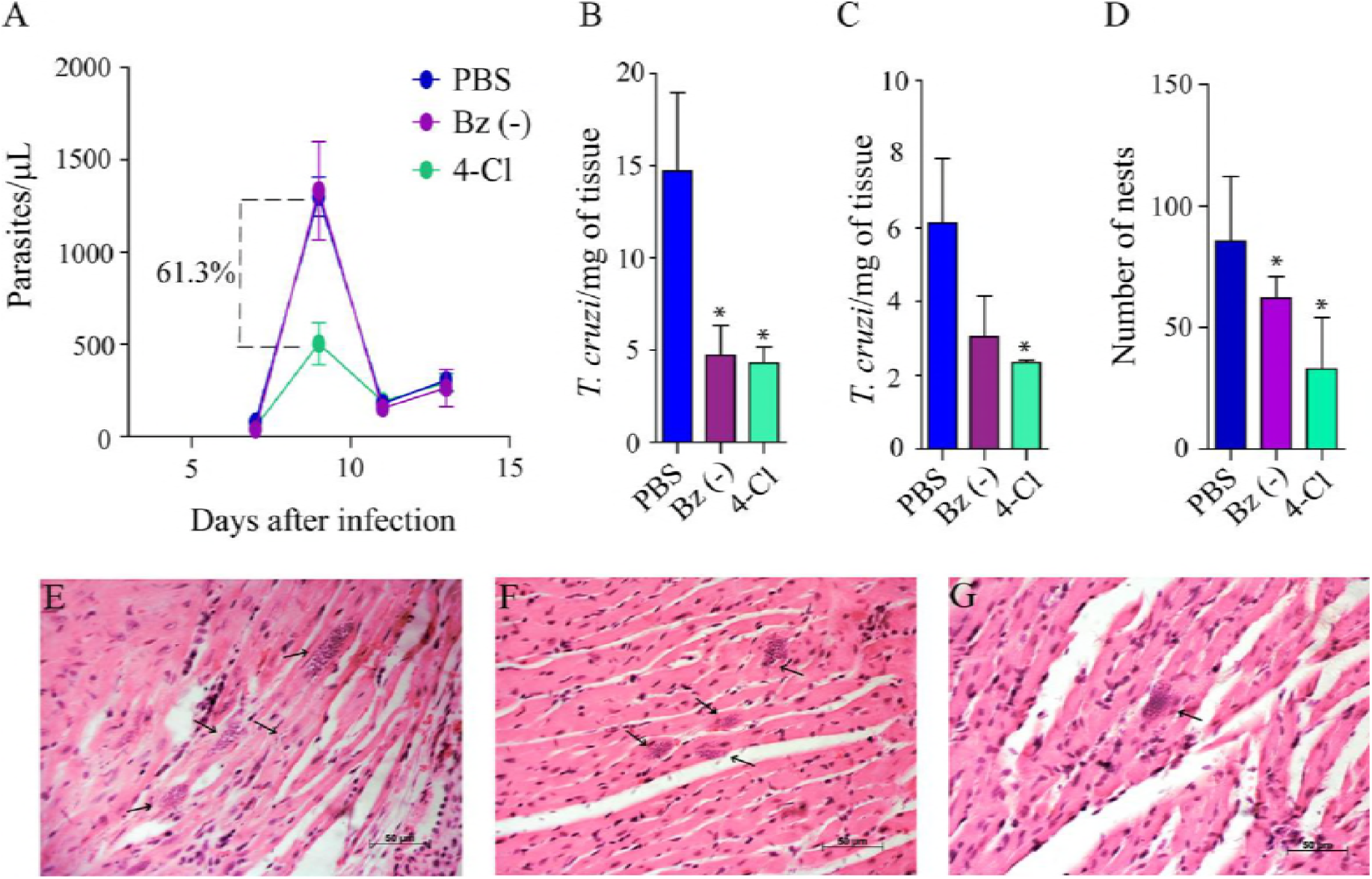
Parasitic burden in the blood and tissue of infected mice after treatment with 2.8 mg/kg/day of 4-Cl, 2.8 mg/kg/day of Bz(-) and 100 mg/kg/day of Bz(+) during the acute phase of infection. (A) Results indicated the decrease in parasitaemia of animals treated with 4-Cl. Parasitaemia was monitored on days 7, 9, 11 and 13 after infection. (B) Quantification of the parasite burden in cardiac tissues via real-time qPCR. The presence of *T. cruzi* in infected heart tissues of mice was analysed by qPCR 15 d.p.i. (C) Quantification of the parasite burden in skeletal tissues via real-time qPCR. The presence of *T. cruzi* in infected skeletal tissues of the mice was analysed by qPCR 15 d.p.i. (D) Number of amastigote nests in heart tissue. (E) Heart histological section of PBS-treated mice, (F) Bz(-)-treated mice and (G), 4-Cl-treated mice. Arrows indicate amastigote nests. The mean±SEM is shown and represents three independent experiments (n=5). Significance was defined when *p≤0.05.

Infection with the Y strain was shown to generate intense inflammatory infiltrates with few amastigote nests, isolates or small groups (19) as observed in the PBS and Bz(-) heart sections (Fig. 3E-F). In 4-Cl treatment, the presence of isolate nests was also accompanied by a strong inflammatory infiltrate (Fig. 3G). When the survival of treated mice was evaluated, we observed that 4-Cl protected the animals from the acute phase of infection, while all mice succumbed to the infection after Bz(-) treatment as well as the PBS treatment. This protection conferred by 4-Cl was the same as that of Bz at its optimised dose (100 mg/kg/day), referred to as Bz(+) (Fig. 4A). After 150 days post-infection, the hearts of the animals treated in the acute phase that survived the infection were removed, and *T. cruzi* DNA was measured. The parasite load were detected, and the levels in the hearts of 4-Cl-treated mice were as low as those in Bz(+)-treated mice (Fig. 4B), although it was not possible to observe either the amastigote nests or the inflammatory infiltrate in the heart of both treated groups (Fig S2). These differences between high parasitaemia and low parasitism in the Bz(-) treatment can indicate that the trypanocidal effects of Bz probably occur throughout the treatment, shortly after the peak of infection, since the mice continued to be treated until they were sacrificed. In the thirteenth d.p.i, the parasite levels in the blood of the Bz(-) group showed a reduction compared to the PBS-treated mice, although this reduction was not statistically significant (Fig. 3A).

**Fig 4.**
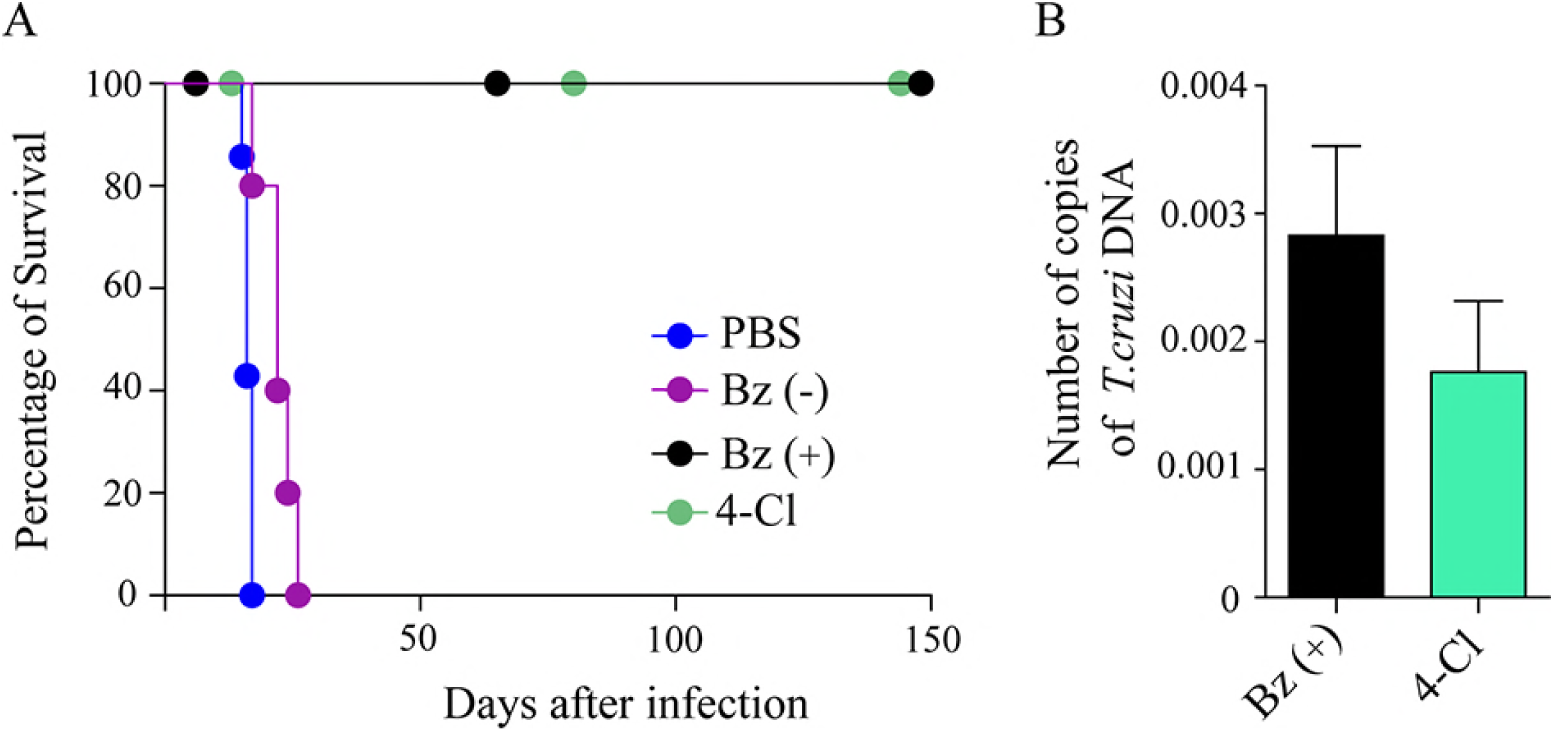
The 4-Cl treatment protects the mice from *T. cruzi* infection. (A) *T. cruzi*-infected mice (n=7) were treated for 10 consecutive days, from the fifth day after infection, with a Bz(-) concentration of 2.8 mg/kg, a Bz(+) concentration of 100 mg/kg (positive control), a 4-Cl concentration of 2.8 mg/kg and PBS. The mice were followed up for 150 days to evaluate survival. (B) *T. cruzi* DNA quantification in the hearts of surviving mice (n=3).

In the chronic phase, the Bz(+)-treated mice tended to show an increased cardiac parasitic burden compared to 4-Cl-treated mice (Fig. 4B). Another point to highlight is the number of parasite nests found in the cardiac tissue of treated mice. In the histopathological sections, Bz(-)-treated mice presented large numbers of nests with a reduced area compared to 4-Cl-treated mice (Fig. 3F and 3G). These smaller nests with greater quantities (Fig. 3D) may have resulted in a similar quantification of parasites in the PCR assay between Bz(+)- and 4-Cl treated mice (Fig. 4B). Together, these data suggest that the 4-Cl treatment in the acute phase protects the mice from lethal *T. cruzi* infection.

### 4-Cl do not show tissue toxicity

Analysis of toxicity is essential for the evaluation and development of new drugs for the treatment of diseases. One way to evaluate the toxicity *in vivo* is to quantify the activity of the enzyme Aspartate Aminotransferase (AST), which is detected in the cytoplasm and mitochondria of a variety of tissues, such as the liver, heart, skeletal muscle, pancreas, and red blood cells, and, therefore, is indicative of systemic damage (17) There was no difference in toxicity between the treatments (Fig S3), revealing that the high levels of AST found may be due to the *T. cruzi* systemic effects (20). However, Alanine Aminotransferase (ALT) is primarily a liver-specific enzyme. The organometallic treatment, similarly to Bz(-), reduced the ALT levels in the serum of infected mice after 15 d.p.i. (Fig 5A). Interesting, this reduction continuous expressive when compared to Bz(+) treated mice after 150 d.p.i (Fig 5B).

**Fig 5.**
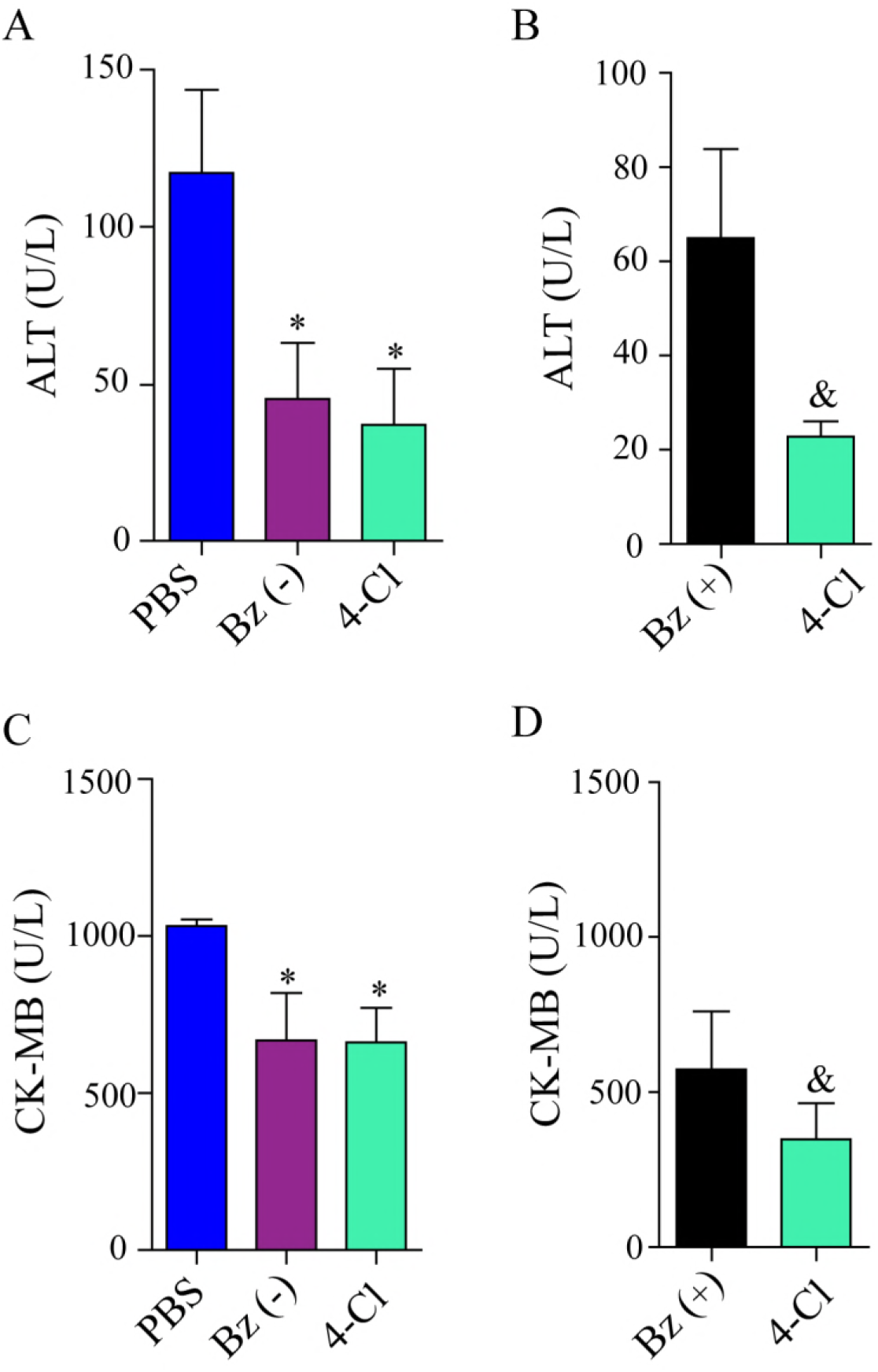
Liver and cardiac lesions of *T. cruzi-*infected mice after treatment with 4- Cl. Quantification of (A) Alanine Aminotransferase (ALT) 15 d.p.i. (acute phase), (B) Alanine Aminotransferase (ALT) 150 d.p.i. (chronic phase), (C) CK-MB (U/L) at 15 d.p.i. and (D) CK-MB (U/L) at 150 d.p.i. The data are represented as the mean ± SEM of three independent experiments, (n=5), using Student’s t test and Mann-Whitney *post test* analysis. Data were considered significant when p<0.05. (*) indicates difference from PBS-treated mice and (&) differences from the Bz(+)-treated group.

In the other hand, skeletal muscle is the target tissue in the *T. cruzi* infection. Serum analysis of 4-Cl infected-treated mice showed a significant reduction of CK-MB, the enzyme released into the plasma during cardiac lesion, in acute phase (Fig 5C) and reduced 3 times the CK-MB levels in the surviving-treated mice (150 d.p.i) as compared to Bz(+) treated mice (Fig 5D). Together, these data reinforce a protective role of 4-Cl treatment.

### High levels of IFNγ are detected in the early stage of acute infection after 4-Cl treatment

Studies suggest that the treatment in cooperation with host immune system has a large impact on the clearance of parasite (21, 22). Th17 cells act early in the infection by releasing IL-17A, which promotes activation of the phagocytosis respiratory burst response and indirect activation of CD8+ T cells(23, 24). Then, we verified the production of IL-17A in the serum of infected mice at both 15 and 150 d.p.i. There was no difference in the production of this cytokine between the treated groups (Fig S4). Although, recent report showed that Th17 are an important impact on protection against *T cruzi* infection, the classical immune protection is based on Th1 response (24). Proinflammatory cytokine production, such IL-12, TNF and IFN-γ are required to activate T lymphocytes, macrophage and other cells, resulting in parasite control (25–27). Surprisingly, we observed an increase of IFN-γ production in 4-Cl-infected treated mice, higher than mice treated wih Bz(-) but lower than those treated with saline (Fig. 6A). This high production of IFN-γ remained constant until chronicity of the disease, at levels similar to the optimal dose of Bz (Bz+) (Fig. 6B). This fact may suggest that the 4-Cl may modulate the immune response at the beginning of infection.

**Fig 6.**
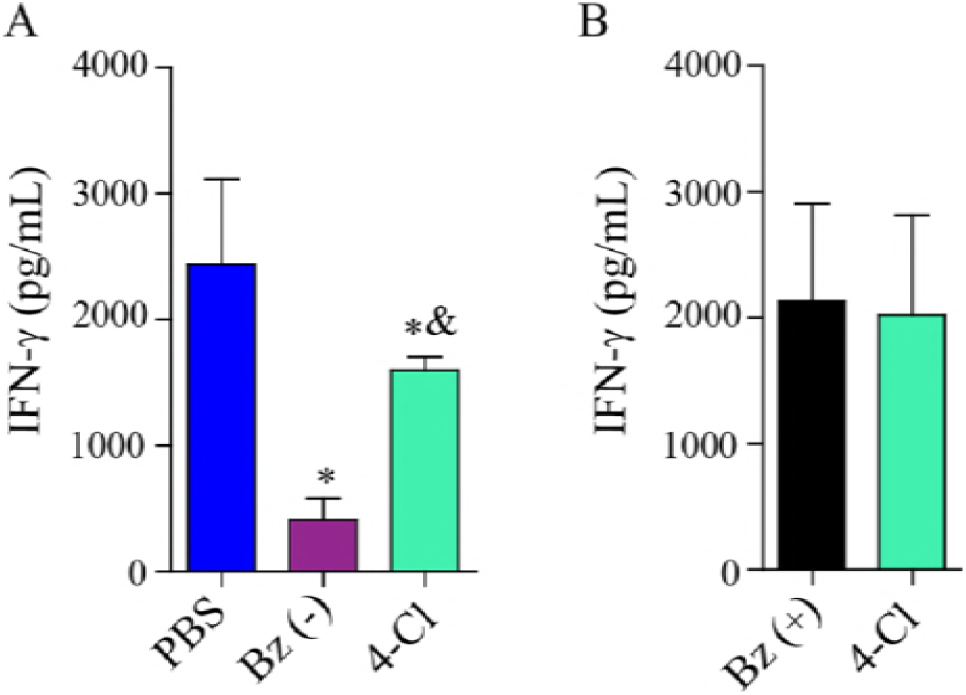
Treatment with 4-Cl increases the systemic production of IFN-γ in animals infected with *T. cruzi*. IFN-γ was measured by ELISA in the serum of Balb/C mice infected and treated with Bz(-) (2.8 mg/kg/day) and/or 4-Cl (2.8 mg/kg/day) and Bz(+) (100 mg/kg/day). (A) Acute phase – 15 d.p.i. (n = 5) and (B) chronic phase – 150 d.p.i. (n = 7). Student’s t test and nonparametric data were compared with the Mann-Whitney U test. Significant differences compared to the control or Bz(-) are denoted by ^&^p<0.05 and ^*^p<0.05, respectively.

### In silico binding study of 4-Cl

The activity of the immune system induced by 4-Cl is substantial; however, 4-Cl is considered a hybrid molecule, and the presence of two or more mechanisms of action for this molecule is not surprising. Therefore, the investigation of other possible targets inside *T. cruzi* is recommended (28). Since thiosemicarbazones are known to inhibit the *T. cruzi* protease cruzain, an enzyme that has essential functions for parasite survival as discussed above, a molecular docking simulation was performed with this enzyme to identify its possible role as a target of 4-Cl in the parasite. The predicted binding mode of the highest ranked compound with cruzain is presented in Fig. 6. It shows the electrostatic (Fig. 7A) and the hydrophobic (Fig. 7B) cruzain surfaces interacting with 4-Cl. The binding pocket core is primarily composed of basic and hydrophobic amino acid residues. The compound from this study strongly interacts with thirteen cruzain hydrophobic residues as shown in Fig. 7C. The binding mode of 4-Cl was compared with those of two other inhibitors whose crystal structures with cruzain were determined by X-ray crystallography and show interactions in the same pocket (29, 30) (Fig S5). Many basic and hydrophobic amino acid residues from cruzain were shown to be important for *in vivo* regulation of cruzain activity (29). No secondary binding site was found by the docking simulations with the studied compound (Fig S6), although two uncharacterised cruzain binding sites were found in a recent theoretical investigation (31).

**Fig 7.**
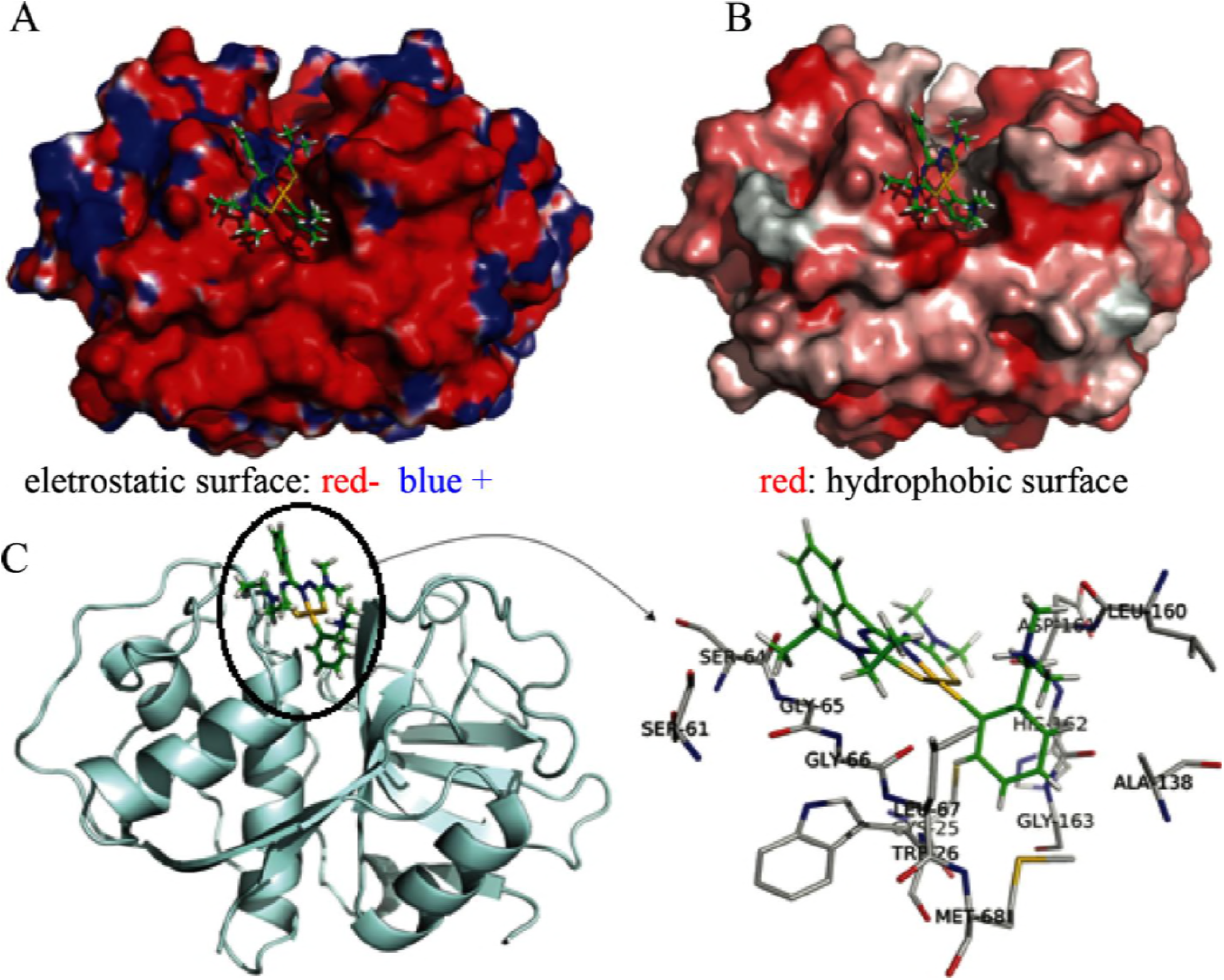
Molecular docking results of the complex formed by the cruzain protease with 4-Cl bound to the enzyme active site. A) Electrostatic surface representation of the complex. B) Hydrophobic surface representation of the complex. C) Cartoon representation of cruzain with the compound in green. D) Hydrophobic residues from the cruzain binding pocket interacting with 4-Cl. Charged residues are coloured blue (positive) or red (negative), and hydrophobicity ranges from high (red) to low (white).

Another notable interaction is with the cysteine sulphur (SG) atom from residue Cys25, which is approximately 3.0 Å away from the S and Au atoms of [Au(Hdamp)(L1^4^)]Cl, as presented in Figure 8. The active site cysteine Cys25 was previously shown to interact via a hydrogen bond with a hydroxymethyl ketone inhibitor and to be a mode of inhibition of cruzain (32). In fact, SG from Cys25 is oriented in the direction of the Au atom from [Au(Hdamp)(L1^4^)]Cl, which suggests formation of a possible covalent bond with the compound. The sulphur atom from the cruzain active site, residue Cys25, was previously reported to be covalently bound to the Z-Phe-Ala-FMK inhibitor (32) and many others inhibitors as well, especially vinyl sulfone derivatives (12, 33, 34). Synthesis and in vitro evaluation of gold(I) thiosemicarbazone complexes for antimalarial activity (34). New experimental data from the crystallization of this compound inside cruzain are necessary to confirm how 4-Cl is bound to cruzain. Although the compound is stable in solution, we cannot determine from the current data if the effect is directly caused by this molecule or its metabolites. In addition, the interaction with other enzymes from *T. cruzi* cannot be eliminated.

**Fig 8.**
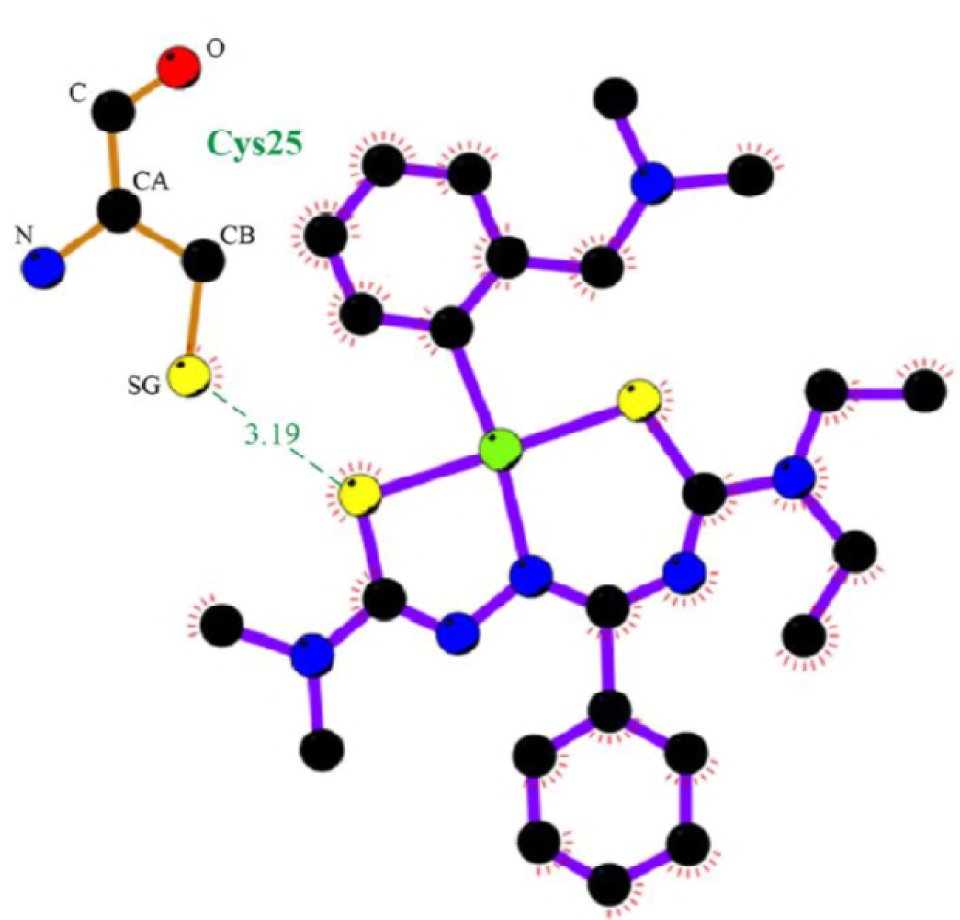
The cruzain active site cysteine Cys25 is predicted to be ~3.0 Å away from the S2 and Au atoms of 4-Cl.

## Discussion

The 4-Cl treatment on *T. cruzi-*infected mice is an interesting treatment because its low concentration, stability and when used, in a very short-term treatment could significantly improve the survival of the animals. Besides that, the synthesis is quite simple, occurring at room temperature in a short period of time, and has an excellent yield (almost 90%). The compound is easily crystallised, which leads to a high purity. Therefore, although it is a gold compound, the costs for preparation do not limit its use. Furthermore, the ESI^+^ MS spectrum shows an exclusive [M]^+^ molecular ion peak with no evidence for the formation of gold(I) compounds, a common finding for gold(III) thiosemicarbazone derivatives. Even when incubated in aqueous solution or in the presence of a reducing agent, such as glutathione, for 24 h at 37 °C, the compound did not show changes in retention time on HPLC (8). This finding is consistent with the high stability of 4-Cl, which is a consequence of the organometallic Hdamp moiety. These characteristics, together with its promising biological activity, are necessary for a drug candidate.

In very low dosage, 4Cl was shown to be significantly selective against both tryposmastigote and amastigote forms of parasite *in vitro* but presented a reduction of nearly 60% of parasitemia, *in vivo*, without alteration in the parasitic load when compared to the same dosage of Bz(-) in the acute phase of infection. This apparent inefficiency in reducing cardiac parasitism, at the onset of infection, may be justified by intrinsic distribution characteristic of the strain. The Y strain is known for its fagotropism at the beginning of infection, reaching organs such as the spleen, liver and bone marrow, and later moves to the muscle cells from the skeletal muscle and heart (35). The quantification of PCR parasitism in the heart at 15 d.p.i. did not reflect the animal′s parasitic burden in the organism. Consequently, it is difficult to conclude that all parasites quantified in the blood were reallocated to the heart. This temporal synchrony became clearer when we observe the cardiac parasitism in the chronic phase in which the 4-Cl treatment was more efficient than the full dosage of Bz (Bz+) in eliminating the parasites of heart demonstrating an intrinsic 4-Cl ability to protect the mice of infection.

One of the aggravating factors of the therapy with Bz are its side effects. At a dose of 100 mg/kg, Bz is hepatotoxic due to high levels of reactive metabolites that are directly generated during Bz metabolism (6, 20, 36). The toxicity of Bz is higher than the damage caused by *T. cruzi* infection. The combination of infection plus Bz treatment by itself increases the levels of both AST and ALT (20). There are no reports showing ALT levels in chronic-phase Bz-treated mice. Our report is the first to demonstrate this small recovery of hepatic lesions after Bz treatment in the acute phase of *T. cruzi* infection. Additionally, Santra et al. (2000) (37) showed that 6-month-old Balb/C mice had baseline levels of 24.30 U/L. These values are very similar to values found for 4-Cl-treated mice after 150 d.p.i. (~20 U/L) that showed a higher reduction of the ALT enzyme than that for the clinical dose of Bz(+) (Fig. 5B), indicating that the 4- Cl treatment is not toxic to the liver. In addition, the toxicity demonstrated during the end of the 4-Cl treatment (50 U/L) was likely due to the parasite burden present in the acute phase of infection. Consistent with these findings, the AST concentration in the serum of C57BL/6 mice, which are resistant to *T. cruzi* infection, is 50 U/L after 15 d.p.i (20), confirming that the little hepatotoxicity found after the treatment may be due to infection and not directly due to some metabolite of the compound.

*T. cruzi* shows preferential tropism for the muscle tissue. Locally, the presence of parasites recruits inflammatory cells to control and eliminate parasitaemia and tissue parasitism, but eventually, this event leads to death caused by *T. cruzi*-induced myocarditis (38). Within the first 15 days of treatment, the reduction of both parasitaemia and cardiac parasitism (Fig. 3A and B) was reflected in the decrease in blood CK-MB levels after 4-Cl treatment (Fig. 5C). However, this reduction was not sufficient to reach the basal levels of uninfected mice, indicating that infection itself or the recruitment of inflammation, for example, may damage cardiomyocytes during the acute phase of the disease. This reduction is substantial over the time, and it was shown to be significant compared with that of Bz(+). Together, these results emphasise the protective effect of 4-Cl during the acute *T. cruzi* infection without causing toxicity to the mammalian organism.

The protection against *T. cruzi* infection is accomplished by an efficient immune response of Th1 profile cells, the major producers of IFN-γ cytokines, and more recently by effector CD4+ T helper lymphocytes, called Th17 cells, which primarily produce interleukin 17 (IL-17). Together, these immune responses are essential for controlling the parasitism and cardiac inflammation during the infection (23). However, excess Th1-biased CD4+ T cells orchestrate a CD8+T cell response that causes tissue destruction and fibrosis (39, 40). Consequently, a balance of this response is crucial for survival of the organism during the infection (39). As shown in this study, the 4-Cl treatment reduced the parasitic burden (Fig. 3), which decreased antigenic exposure and reduced the recruitment of immune cells (41). Therefore, we expected a reduction of IFN-γ production in the serum of treated mice 15 d.p.i. Surprisingly, the gold(III) complex increased the IFN-γ production compared to that in Bz(+)-treated mice, but it was significantly reduced compared with that of the saline group (Fig. 6A) at the early stage of infection. In accordance with our data, during the acute phase, the intense parasite burden activates the murine immune system to produce approximately 2000 pg/μL of IFN-γ in the Balb/C (23) serum and 200 pg/μL in C57BL/6 mice (20, 42). Novaes et al. (20) showed that after Bz treatment, the IFN-γ levels were reduced by half in C57BL/6 mice. Therefore, a reduction in the levels of this cytokine is expected after Bz treatment. The acute phase of *T. cruzi* infection is marked by an intense inflammatory infiltrate that is reduced when the disease reaches the chronic stage.

Throughout the infection, the cytokine levels were reduced to normal levels comparable to those of the Bz(+) group (Fig. 6B). At this late stage, there is a significant amount of CD8 T cells producing IFN-γ (43) and perforin (44), which are responsible for the maintenance of high levels of this cytokine (44, 45), while secreted perforin leads to late lesions in muscle fibres and cardiac alterations (44). Although treatment with Bz reduces the parasitic burden and, consequently, the inflammatory infiltrate, the action of Bz remains still dependent of the amount of IFN-γ produced. When IFN-γ knockout mice were treated with Bz, they were more susceptible than those treated with posoconazole (46). Our results contrasted to the amount of IFN-γ induced by Bz (Fig. 6). The IFN-γ production was high in the acute phase and decreased in the chronic stage. The early increase of this cytokine may be important to combat the parasites during the acute phase, since IFN-γ acts directly on macrophages, increasing their trypanocidal capacity (47). However, this change may also damage the cardiac tissue, as shown by the elevated levels of CK-MB in the serum of infected and 4-Cl-treated mice. In addition, the reduction of the parasite nests in the cardiac tissue (Fig. 3D) should be reflected in the reduction of IFN-γ production at the chronic stage of 4-Cl-treated mice. The high levels of IFN-γ at the beginning of the infection may be protective if combined with the other parameters, such as reduction of the parasitic burden, cardiac and hepatic protection and mouse survival as long as that in the Bz(+) group. However, it may also damage the cardiac tissue, as shown by the elevated CK-MB in the serum of infected and 4-Cl-treated mice. Together, these results suggest that treatment with 4-Cl may act in the immune system by promoting the release of IFN-γ, which acts directly on parasite death.

Similarly, treatment with an equal concentration of Bz was not able to produce these improvements. As demonstrated by our in silico data, the complex gold(III)- thiosemicarbazone probably interacts with cruzain, a major cysteine protease of *T. cruzi* crucial to multiplication and amastigote survival of *T. cruzi* (8, 10), which can be shown by its direct action on the amastigote and trypomastigote forms of *T. cruzi* in both the Y strain and the Tulahuen strain (8). In addition, the lower levels of tissue parasitism in both skeletal muscle and heart of 4-Cl-treated mice were followed by a reduction of cardiac and hepatic tissue damage, showing a protective effect of this compound. Curiously, 4-Cl could increase the production of systemic IFN-γ, which indirectly kills the parasite by the recruitment of an efficient immune response against *T. cruzi* (48). Therefore, the gold(III) complex appears to act on two different pathways of parasite death. As predicted by our previous study (8), this complex may have multiple targets and mechanisms of action by affecting different biological functions in the parasite. Future studies will reveal how the 4-Cl complex modulates the immune response: whether the increase of IFN-γ can activate macrophages to produce nitric oxide, the most classic form of *T. cruzi* death by immune system (40); if it inhibits other parasite enzymes, such as TcOYE (49, 50); or if this complex produces intracellular reactive oxygen species, which are known to cause damage to *T. cruzi*. In any case, 4-Cl was shown to be a potential candidate for the treatment of Chagas disease.

## Conclusion

The gold(III) complex 4-Cl was shown to be effective in directly killing the parasite in both the trypomastigote and amastigote forms of *T. cruzi.* The *in vivo* assays demonstrated that 4-Cl reduces parasitaemia and tissue parasitism at a very low dose in addition to protecting the liver and heart from tissue damage, leading to the survival of 100% of the 4-Cl-treated mice during the acute phase. The same effect has been observed only when the mice were treated with the maximum dose of Bz. During the acute phase, we observed a specific increase of systemic IFN-γ, a classical cytokine for protection of the organism during Chagas disease. Interestingly, during the chronic phase, the production of this cytokine returns to the same levels as that of the Bz(+) treatment, which indicates that 4-Cl can modulate the immune response during the treatment period. Theoretical studies (molecular docking) suggest that 4-Cl could interact with cruzain via hydrophobic interactions and that it might react with Cys25, a residue commonly found to be covalently bound to many other cruzain inhibitors. Future studies on the biochemical and metabolic pathways are necessary to further elucidate the anti-*T. cruzi* mechanism of action of this drug candidate. However, from the data presented thus far, 4-Cl appears to act on two different pathways. This study has confirmed that the organometallic gold(III)-thiosemicarbazone complex 4-Cl may be a new candidate for the development of novel anti-chagasic drugs and may accelerate the investigations based on metallotherapeutics for the treatment of Chagas disease.

## Materials and Methods

### Reagents and supply

The synthesis of 4-Cl has been described previously (8). Benznidazole was purchased from Sigma-Aldrich (used as a reference drug). RPMI medium 1640 (with or without phenol red) supplemented with 5% bovine fetal serum (GIBCO, Grand Island, NY, USA), 100 IU mL^−1^ penicillin G, and 100 mg mL^−1^ streptomycin (Gibco-BRL, Grand Island, NY, USA) was used. Dimethyl sulfoxide (DMSO) was obtained from Sigma-Aldrich Chemicals, Co. (St. Louis, MO, USA).

### *T. cruzi* stocks

All the procedures and animal protocols were conducted in accordance with the National Brazilian College of Animal Experimentation (COBEA) and approved by the Commission of Ethics in Animal Research of the University of São Paulo, Medical School of Ribeirão Preto (CETEA) - Protocol number 100/2014.

For the *in vitro* experiments, the LLC-MK2 cells were infected with bloodstream trypomastigote forms, which were derived from previously infected Swiss mice. For the *in vivo* experiments, mice were intraperitoneally inoculated with 2000 bloodstream trypomastigote forms, also derived from previously infected Swiss mice.

### *In vitro* evaluation of trypanocidal activity against the trypomastigote and amastigote forms

Trypomastigote forms of the *T. cruzi* Y strain were obtained from infected LLC-MK2 cell culture and suspended at a concentration of 6.5×10^6^ parasites/mL in RPMI 1640 liquid medium without phenol red. They were cultured in flat-bottom 96-well plates at various concentrations of Bz or 4-Cl at 37 °C for 24 h. The viability of the parasites was determined by counting the motile parasites in a Neubauer chamber as previously described (7, 51). The concentration of the compound corresponding to 50% trypanocidal activity in trypomastigote was expressed as the IC_50Try_.

For evaluation of the trypanocidal activity of the compound against the amastigote forms of the *T. cruzi* Y strain, differentiated bone marrow macrophages (BBMs) were used as previously described (52), and femurs were obtained from 6-8-week-old C57BL/6 mice. Cells were seeded in non-tissue culture-treated Optilux Petri dishes (BD Biosciences) and incubated at 37 °C in a 5% CO_2_ atmosphere. Four days after seeding the cells, an extra 10 mL of fresh R20/30 was added per plate and incubated for an additional 3 days. The supernatants were discarded, and the attached cells were washed with 10 mL of sterile PBS to obtain the BBMs. The macrophages were detached by gently pipetting the PBS. The cells were centrifuged at 200× g for 5 minutes and resuspended in 10 mL of BBM cultivation media (R10/5). The cells were counted, seeded (5.0×10^4^ cells/well) and cultivated in tissue culture plates for 24 hours. After 24 hours, the BBMs were infected with trypomastigote forms (1×10^5^ parasites/well) for 16 h. The cells were washed to remove parasites in the supernatant and incubated with Bz or 4-Cl for an additional 24 h at 37 °C. Cells were stained with Giemsa dye and evaluated by optical microscopy [21]. Trypanocidal activity was determined by counting the parasites/cell in at least 200 cells.

### *In vitro* evaluation of the cytotoxicity in BBMs

BBMs were assessed by the colorimetric method [3-4,5-dimethylthiazol-2-yl) 2,5- diphenyl tetrazoyl bromide] (MTT) (53). BBMs were suspended at a concentration of 5.0×10^5^ cells/mL in RPMI medium without phenol red supplement with 5% fetal bovine and incubated for 24 hours in 96-well cell culture plates. After this incubation period, the Bz and Gold(III) complex were added at serially diluted concentrations ranging from 125.0 to 1.95 μM. The cells were incubated for 24 hours at 37 °C. After incubation, the medium was removed, and fresh culture medium containing 50 μM of MTT (2.5 mg mL^-1^) diluted in phosphate buffered (PBS) was added. The precipitated blue MTT formazan was dissolved in 50 μL of DMSO, and the absorbance was measured at 570 nm in a VARIAN CARY-50 multiwell MPR plate reader. Cell viability was expressed as the percentage of absorption values in treated cells compared with untreated (control) cells. The concentration of the compound corresponding to 50% cytotoxicity in the BBMs was expressed as the CC_50BBMs_.

### Animal inoculum and treatment

Balb/c female mice (4 – 6 weeks old) with an average weight of approximately 20 ± 3 g were used to determine the trypanocidal activity of the compounds in the acute phase of Chagas disease. All animals were kept under the same conditions, receiving water and food ad libitum. Animals were infected intraperitoneally with 2.0 × 10^3^ blood trypomastigote forms of the *T. cruzi* Y strain (7, 51). The mice were orally treated, and treatment started at five days post-infection (d.p.i) for 10 consecutive days. The daily doses of the compounds tested were 2.8 mg/kg/day of 4-Cl, 2.8 mg/kg/day of Bz(-), and 100 mg/kg/day of Bz(+). Bz(+) was used as a positive control in the study (7). For analysis of the acute stage of infection, the animals were euthanised after 15 d.p.i. The mice that survived the acute phase were followed for 150 d.p.i., and their organs and serum were removed and stored at −80° C for further analysis.

### Parasitaemia and survival assessment

Parasitaemia was analysed on alternate days from the 5^th^ d.p.i. To this end, 5 μL of fresh blood was collected from the animal tail. The count in 100 fields was performed via direct observation under a light microscope (54). The survival was determined by daily inspection for more than 150 consecutive days in which mice were weighed to monitor the effects of the infection.

### Quantification of cardiac and hepatic injury

The blood of infected mice was collected by cardiac puncture and centrifuged for 10 minutes at 12000 g. Then, the serum was removed and stored at minus 4 °C until performance of biochemical assays. The cardiac lesions of mice infected with *T. cruzi*, treated or not, were assessed by measuring the serum creatine kinase-MB (CK-MB) levels at 15 and 150 d.p.i. The CK-MB levels were measured using a CK-MB kit (LABTEST, Brazil), as previously described (55). Absorbance was measured on a microplate spectrophotometer (EMAX Molecular Devices Corporation, California, EUA). The colour produced from this reaction was measured at a wavelength of 340 nm; the results are expressed in U/L. The hepatic damage was evaluated by measuring Aspartate aminotransferase (AST) and alanine aminotransferase (ALT) using a LABTEST system (Brazil), according to the manufacturer’s instructions. The colour produced by this reaction was measured at a wavelength of 340 nm. The results are expressed in U/L.

### Histological analysis

Groups of five mice were euthanised at 15 and 150 d.p.i., and the hearts were fixed in paraffin for histological analysis. For analysis of amastigote nests via light optical microscopy (Axioskop 40), tissues were sectioned at a 5-μm thickness, stained with haematoxylin-eosin (H&E) and examined under a light microscope with 40 times magnification. Each tissue section was imaged 10 times and used to analyse the amastigote nests.

### Quantitative real-time polymerase chain reaction (qPCR)

The qPCR technique was used to determine the amount of parasite DNA in the tissues collected from infected and treated animals. The DNA was purified from 10 mg of heart tissue using Wizard^®^ Genomic DNA Purification (Promega), according to the manufacturer’s instructions. Each qPCR reaction contains 20 ng of genomic DNA, 0.3 μM of the specific primers of *T. cruzi* (TCZ-F 5′-GCTCTTGCCCACAMGGGTGC-3′ (M= A or C) and TCZ-R 5′-CCAAGCAGCGGATAGTTCAGG-3′’(56)), which amplify a 182 bp product, 7.6 μL GoTaq qPCR Master Mix^®^ (Promega), and H_2_O to a final total volume of 15 μL. Reactions were performed using the StepOnePLus™ Real-Time qPCR System (Applied Biosystems, Foster City, CA, USA). The cycle programme was 95 °C for 10 minutes, followed by 50 cycles of three steps, denaturation at 95 °C for 15 seconds, annealing at 55 °C for 30 seconds, and amplification at 72 °C for 15 seconds. The melting phase was performed at 95 °C for 15 seconds and 60 °C for 60 seconds, followed by a 0.3 °C ramp and 95 °C for 15 seconds. During the melting phase, the acquisition setting was set at step. The data were analysed with StepOne Software version 2.2.2.

### Cytokine quantification by ELISA

For analysis of cytokine production in the serum of treated-infected mice, blood was collected by cardiac puncture and centrifugated at 12000 g for 10 minutes. The serum was collected and frozen at-20 °C. The ELISA sets were IL-17A and IFN-γ (R&D, Minneapolis, MN, USA). All technical procedures were performed according to the manufacturer′s instruction. The limits of sensitivity for different assays were as follows: 15 pg/mL for IL-17A and 31.2 pg/mL for IFN-*γ*.

### Docking studies

The X-ray crystallographic data of the cruzain protease was extracted from the Protein Data Bank (18) (PDB, code 1AIM). Molecular docking simulations were performed with Genetic Optimization for Ligand Docking (GOLD) suite version 5.5. The complete protocol is presented in the **Supporting Information** section.

### Statistical analysis

The statistical analyses are representative of the mean ± SEM of three independent experiments, (n=5), using Student’s t test and Mann-Whitney *post test* analysis. Data were considered significant when p<0.05. (*) indicates differences compared to the PBS-treated mice and (&) differences from the Bz(+)-treated group. All analyses were performed using PRISM 5.0 software (GraphPad, San Diego, CA, US).

## Acknowledgements

The authors gratefully acknowledge the technical assistance of Cristiane Maria Milanezi and Wander Ribeiro. We also thank Rubilan Carneiro Quionero for supervising the animal facility as well as providing all animals used in this study. The research leading to these results received funding from CAPES and CRID, Center for Research in Inflammatory Disease.

## Supplementary Information

**FIG S1.**
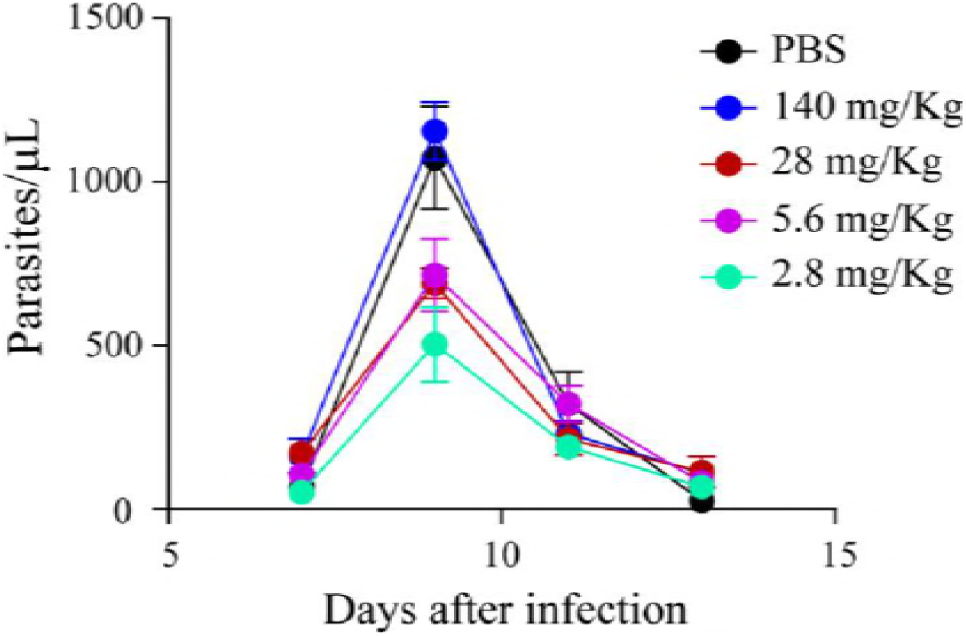
Parasitaemia of mice treated with different concentrations of 4-Cl. The mice (n=5) were treated for 10 consecutive days, starting at 5 days after infection. The choice of concentrations was based on the conversion in molarity of the standard dose of Bz (100 mg/kg). Thus, 140.0 mg refers to a dose five times, 28.0 mg ten times, 5.60 fifty times, and 2.80 a hundred times lower than Bz. The parasitaemia was monitored on days 7, 9, 11 and 13 after infection.

**FIG S2.**
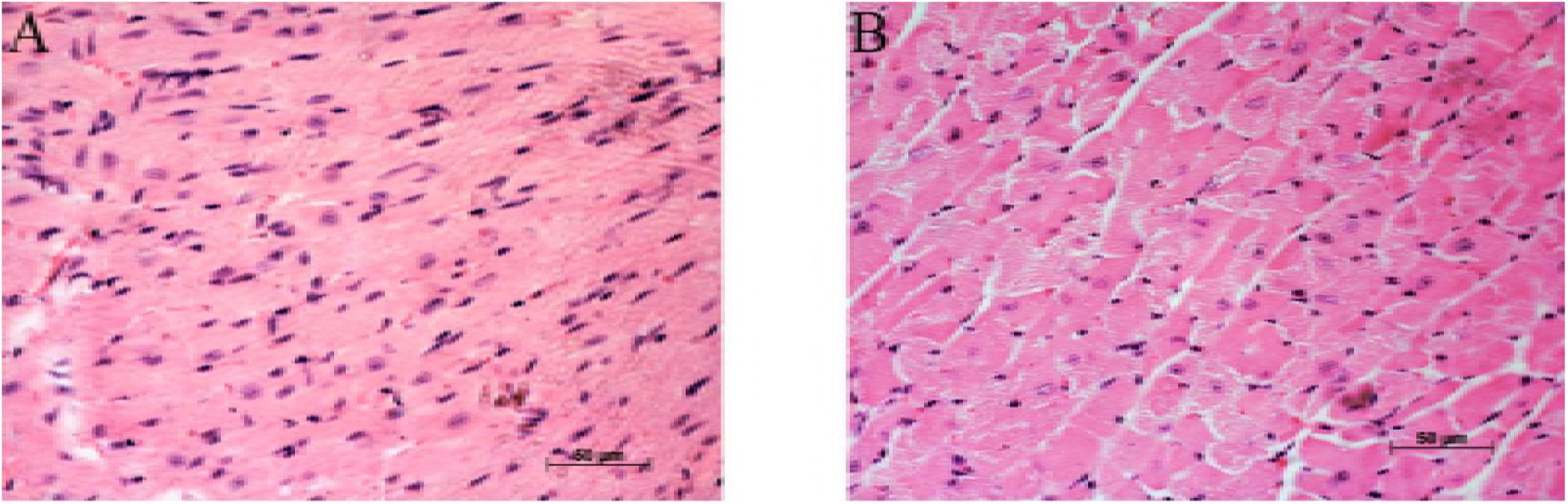
Heart histological sections of treated and surviving mice 150 days after infection. (A) Bz(+) treated-mice; (B) 4-Cl-treated mice.

**FIG S3.**
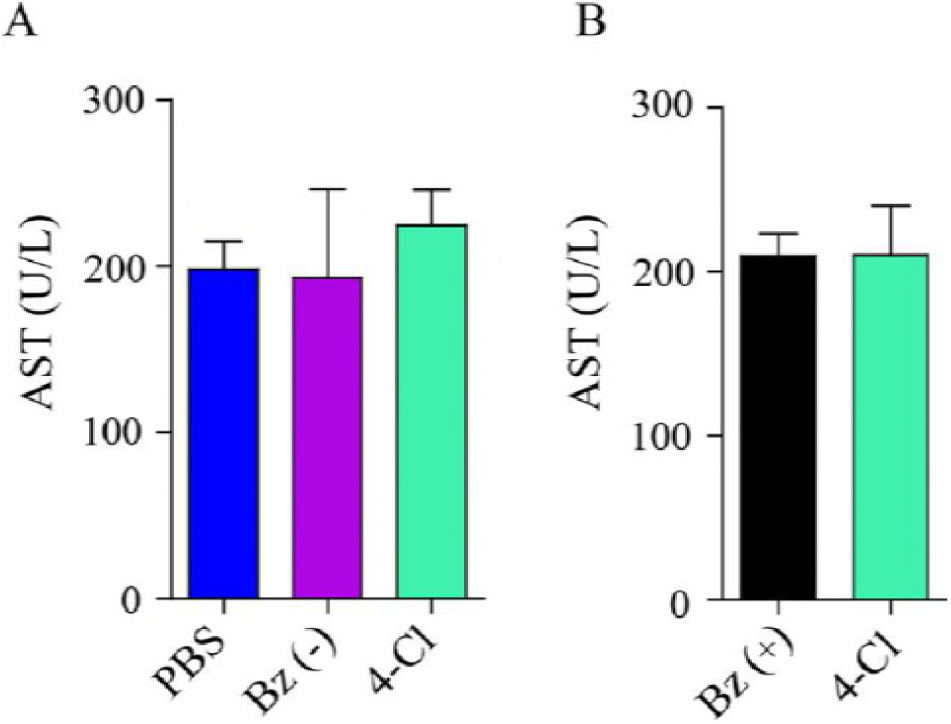
Liver lesions of *T. cruzi-*infected mice after treatment with 4-Cl. (A) Quantification of Aspartate aminotransferase (AST) levels 15 d.p.i. (acute phase, n = 5). (B) Quantification of Aspartate aminotransferase (AST) levels 150 d.p.i. (chronic phase, n = 7).

**FIG S4.**
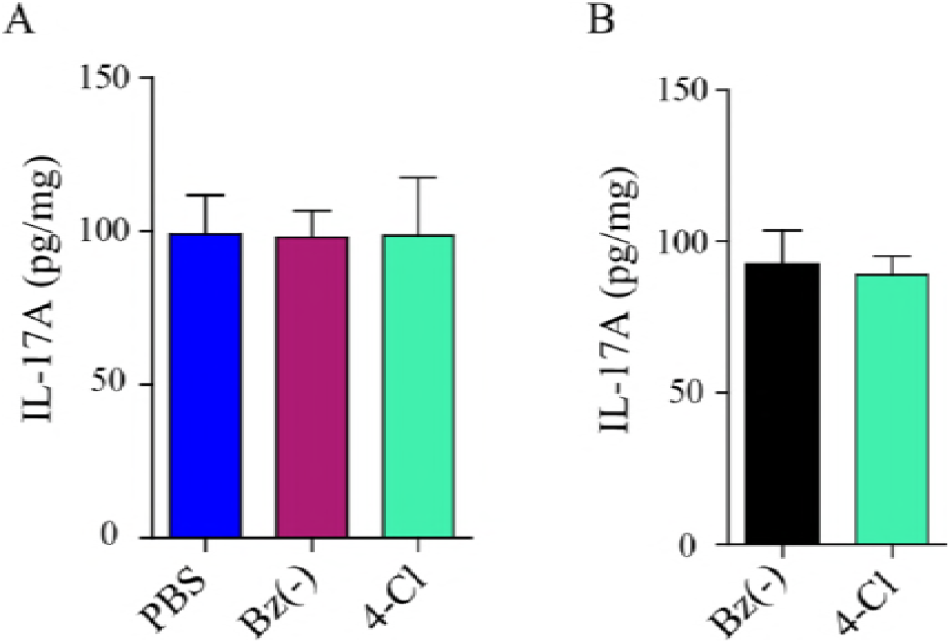
Treatment with 4-Cl did not increase the systemic production of IL-17 in animals infected with *T. cruzi*. IL-17 was measured by ELISA in the serum of Balb/C mice infected and treated with 2.8 mg of Bz and/or 4-Cl. (A) Acute phase - 15 d.p.i. (n = 5) and (B) chronic phase – 150 d.p.i. (n = 7).

**FIG S5.**
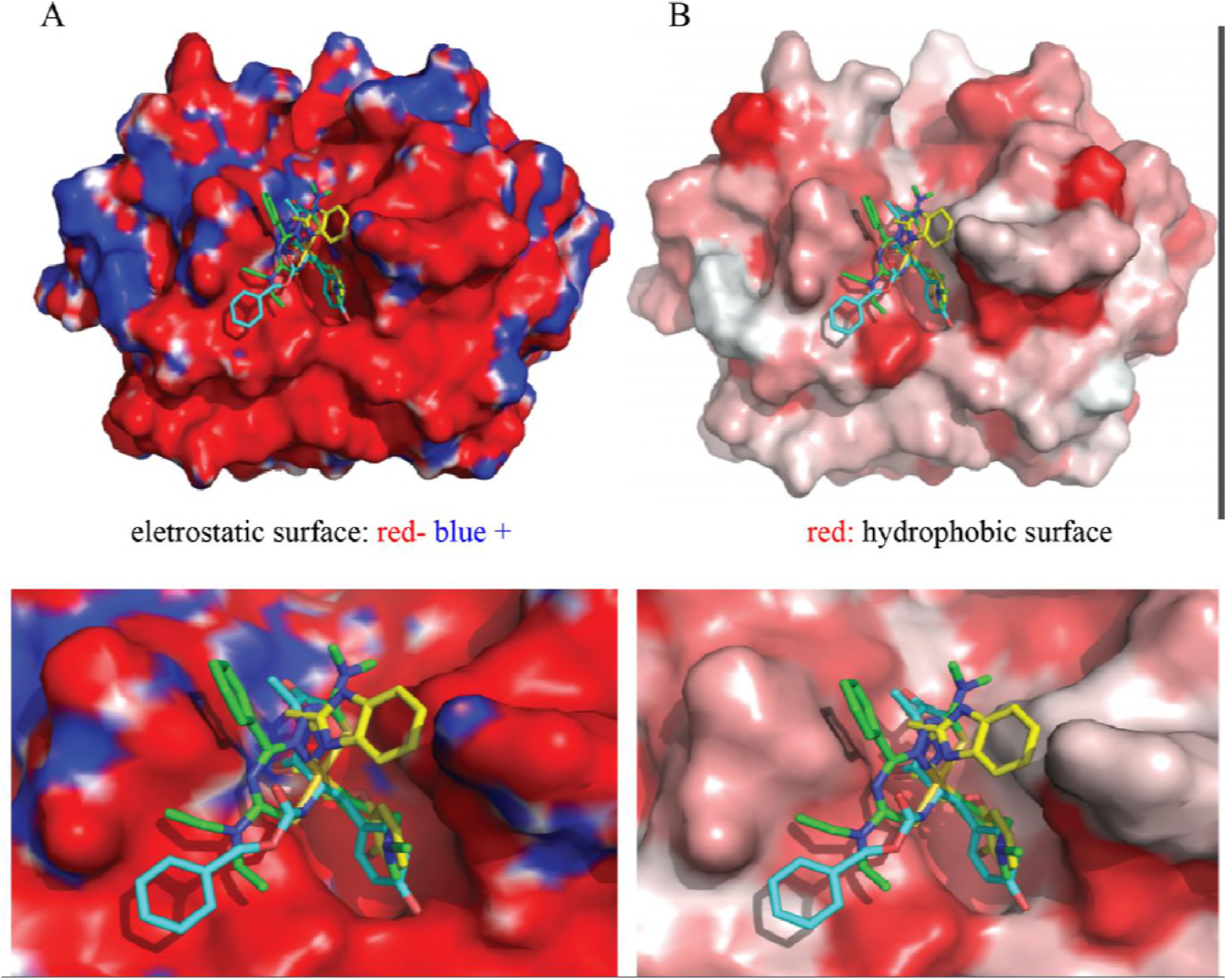
A) Electrostatic and B) hydrophobic surfaces of the cruzain protease with the following compounds: 4-Cl from this study (green), benzoyl-tyrosine-alanine-fluoromethylketone from the X-ray crystallographic data of cruzain protein used in this study (PDB code 1AIM (31), cyan), benzimidazolethyl-bromophenoxy-acetamide from the X-ray data of cruzain protein used in a high-throughput screen of cruzain inhibitors (PDB code 3KKU (30), yellow).

**FIG S6.**
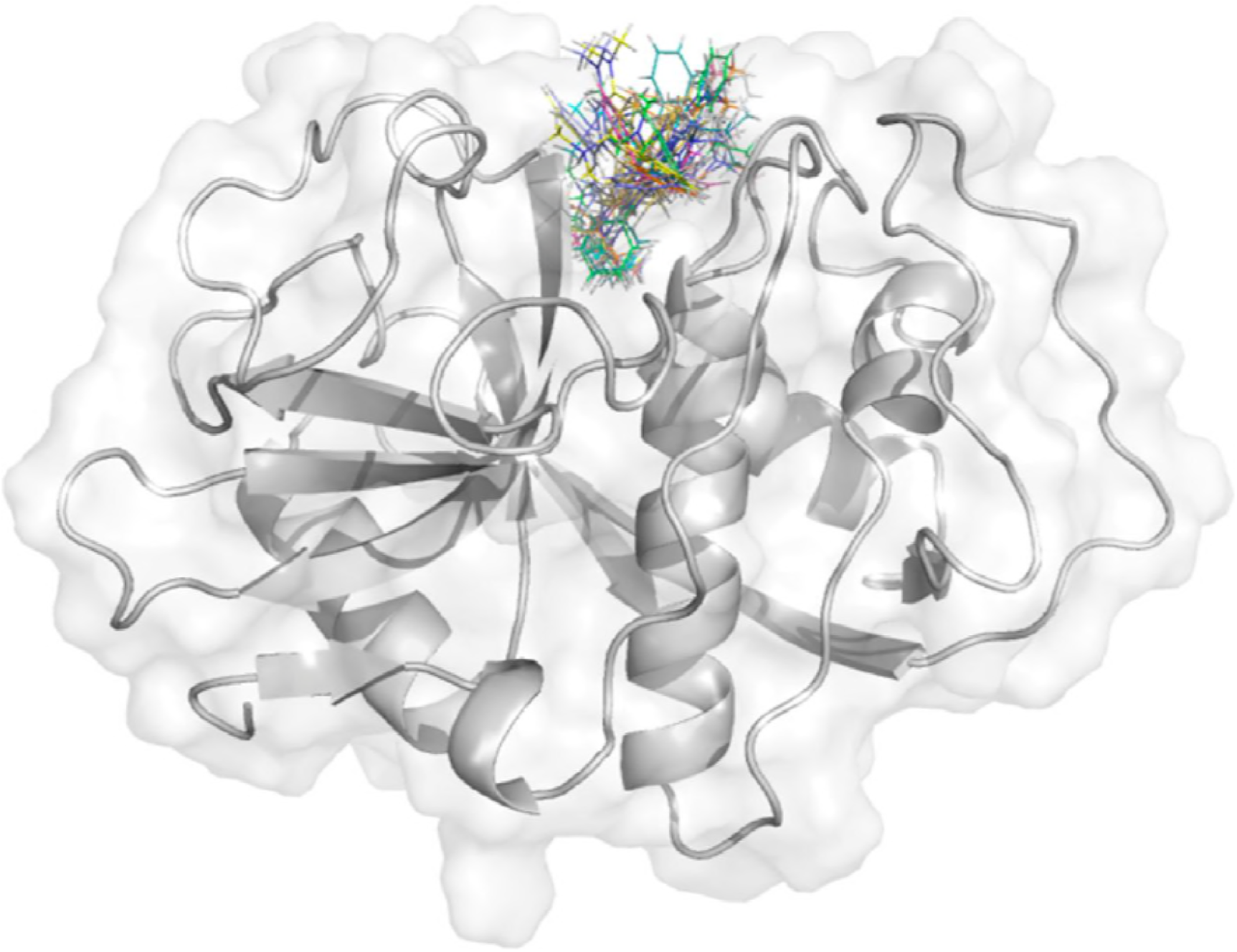
Top 10 best docking solutions of the complex formed by cruzain with [Au(Hdamp)(L)]Cl. All solutions were docked in the primary pocket, the active binding site (29, 30). Compound conformations are shown in lines, and the enzyme is shown in cartoon and surface representations.

## Docking protocol

Molecular docking simulations of the 4-Cl ligand were carried out to explore the binding modes in the cysteine protease cruzain. The X-ray crystallographic data of the cruzain protease were extracted from the Protein Data Bank (57) (PDB, code 1AIM (31). The compound structure was obtained by single-crystal X-ray studies and prepared with GaussianView 6 software (Gaussian, Inc.) and Marvin Sketch suite version 17.1.27, ChemAxon (http://www.chemaxon.com). The docking simulations were performed with the GOLD suite version 5.5 in the enzyme active site. Pre- and post-docking visualisation and interactive docking setup were conduct with Hermes software from the GOLD suite. All water molecules and heteroatoms were removed from the protease. Only residues within 6.0 Å surrounding the native inhibitor were used as the ligand cavity site. The Genetic Algorithm (GA) method was used to run the calculations (30). Full flexibility was allowed to the ligand. GA runs conducted herein with a maximum of 100,000 GA operations were performed on a population size of 100 individuals. Diverse solutions were generated, ring corners were allowed to flip, conformations were explored, and no constraint was applied to the protein or to the ligands. Redocking simulations of the native inhibitor from the PDB code 1AIM were executed in the enzyme active site with all the available GOLD score functions, and the best score (lowest rmsd to the X-ray conformation) was found with the GoldScore fitness function rescored by ChemScore (49). Thus, GoldScore rescored by the ChemScore fitness function was chosen to predict the binding mode of the compound in the study with the cruzain crystallographic structure. The GoldScore ranking was evaluated and the best pose, the highest-ranking structure for the compound, was chosen for the interaction analysis. Figures of the cruzain protease/inhibitor complexes were prepared with PyMOL (Molecular Graphics System, version 1.8 Schrödinger, LLC).

